# Interferon-γ signaling in eosinophilic esophagitis has implications for epithelial barrier function and programmed cell death

**DOI:** 10.1101/2024.01.26.577407

**Authors:** Megha Lal, Caitlin M. Burk, Ravi Gautam, Zoe Mrozek, Tina Trachsel, Jarad Beers, Margaret C. Carroll, Duncan M. Morgan, Amanda B. Muir, Wayne G. Shreffler, Melanie A. Ruffner

## Abstract

**Objective:** Eosinophilic esophagitis (EoE) is a chronic esophageal inflammatory disorder characterized by eosinophil-rich mucosal inflammation and tissue remodeling. Transcriptional profiling of esophageal biopsies has previously revealed upregulation of type I and II interferon (IFN) response genes. We aim to unravel interactions between immune and epithelial cells and examine functional significance in esophageal epithelial cells.

Design

We investigated epithelial gene expression from EoE patients using single-cell RNA sequencing and a confirmatory bulk RNA-sequencing experiment of isolated epithelial cells. The functional impact of interferon signaling on epithelial cells was investigated using *in vitro* organoid models.

**Results:** We observe upregulation of interferon response signature genes (ISGs) in the esophageal epithelium during active EoE compared to other cell types, single-cell data, and pathway analyses, identified upregulation in ISGs in epithelial cells isolated from EoE patients. Using an esophageal organoid and air-liquid interface models, we demonstrate that IFN-γ stimulation triggered disruption of esophageal epithelial differentiation, barrier integrity, and induced apoptosis via caspase upregulation. We show that an increase in cleaved caspase-3 is seen in EoE tissue and identify interferon gamma (IFNG) expression predominantly in a cluster of majority-CD8+ T cells with high expression of *CD69* and *FOS*.

**Conclusion:** These findings offer insight into the interplay between immune and epithelial cells in EoE. Our data illustrate the relevance of several IFN-γ-mediated mechanisms on epithelial function in the esophagus, which have the potential to impact epithelial function during inflammatory conditions.

**Key Messages:** **What is already known about this topic:**

- The transcriptome of esophageal biopsy tissue reproducibly distinguishes eosinophilic esophagitis from histologically normal tissue, with evidence of mixed inflammatory signals.
- Interferon response signature genes are elevated in EoE biopsy tissue compared to controls, suggesting T1 in addition to T2 cytokine signaling within EoE mucosa.

**What this study adds:**

- We observe reproducible, robust upregulation of interferon signature genes in esophageal epithelium, and we confirm that esophageal epithelium expresses functional IFN-a and IFN-γ receptors.
- IFN-γ treatment of epithelial organoids has several detrimental effects, including decreased proliferation, organoid formation, and increased caspase activation.
- Analysis of single-cell RNA-sequencing data from of EoE biopsy tissue during active disease and remission identified that a CD8+ population expressing high levels of *FOS, ITGAE, and ITGA1* expresses high levels of *IFNG*

**How this study might affect research, practice, or policy:**

- We identify esophageal epithelium as the cellular source for interferon response gene signature in EoE and a CD8+ tissue-resident memory T-cell population was the main source of *IFNG*.
- Further mechanistic studies are required to identify how non-T2 signaling mechanisms like IFN-γ signaling contribute to EoE pathogenesis, and if this pathway can be targeted as an adjunctive therapy for EoE.

## INTRODUCTION

EoE is a chronic allergic disorder of the esophagus characterized by eosinophil-rich mucosal inflammation and epithelial barrier dysfunction (1–5). Strong evidence links type 2 (T2) cytokine signaling to mucosal inflammation in EoE, including the presence of pathologic effector T cells in esophageal mucosal during active disease (6–9). The recent successful clinical trials of dupilumab, a monoclonal antibody targeting the IL4Rα subunit of IL-4 and IL-13 receptors, highlight the importance of T2 signaling in EoE (3).

Despite the strong link between T2-cytokine-mediated inflammation and EoE, prior research indicates that IFN signaling is upregulated during EoE(10, 11). Sayej *et al.* measured increased IFN-γ production from EoE biopsy cultures (11). We previously reported that interferon gene signatures (IGS) are upregulated in biopsy tissue from adults and children with EoE (10). This was recently replicated in a large reanalysis of 137 EoE patient biopsy transcriptomes (12).

Here, we utilize complementary datasets to interrogate IFN signaling in EoE. We use single-cell RNA sequencing to examine IFN production and responses in esophageal mucosal cell populations. Within this dataset, we observe heterogeneity of ISG expression, and identify upregulation of ISG within epithelial cells compared to other cell clusters. We confirm the upregulation of ISG in RNA-sequencing of epithelial cells isolated from esophageal biopsies of EoE patients. IFN-γ signaling has previously been shown to decrease epithelial barrier function and induce apoptosis in epithelial cells of the airway, intestine, and skin (13–17). Here, we hypothesized that IFN-γ signaling would result in similar effects in esophageal epithelium and utilized organoid models to examine organoid morphology, epithelial function, and transcriptional responses following interferon treatment. This work offers insight into immune-epithelial crosstalk in EoE and demonstrates potential mechanisms of IFN-γ-mediated esophageal epithelial barrier dysfunction.

## METHODS

### Biopsy single-cell RNA-sequencing

Data from Morgan *et al*. (8) was accessed in NCBI GEO database (8, 18, 19) using accession number <u>GSE175930</u>, and analyzed in Seurat using R as described (8, 20–23) The dataset includes 14,242 cells (10,049 active, 4,193 remission) with over 500 unique genes. T/NK cell dataset included 4423 cells with 900 unique genes each (3066 active, 1409 remission). After normalization, variable gene selection and scaling, UMI and percent mitochondrial genes were regressed prior to principal component analysis and UMAP visualization using the top 10 components. Clusters from Morgan *et al.* were used for further analysis (8) to examine interferon response signature genes previously identified in patients with EoE (10). Differential expression was evaluated (FindMarkers), reporting average log2-fold change and adjusted p-values (Bonferroni corrected). Dot and volcano plots were generated with the R packages, dittoSeq 1.11.0 (24) and EnhancedVolcano 1.16.0 (25), respectively.

### Esophageal biopsy specimens

The studies were approved by the institutional review board of Partners Healthcare (GSE175930) and Children’s Hospital of Philadelphia (GSE234973). Participants or their legal guardians provided written informed consent prior to study participation.

Esophageal biopsies for the esophageal epithelial whole sample RNA-seq study were collected during routine clinical endoscopy (Table S1). Subjects met EoE diagnostic criteria (26), with active EoE defined by symptoms of esophageal dysfunction and ≥15 eosinophils/high-power microscopic field on biopsy tissue. Control subjects underwent endoscopy due to clinical symptoms, but esophageal and distal histology demonstrated no abnormalities. Patients with celiac disease, inflammatory bowel disease, gastrointestinal bleeding, immunodeficiency, or recent immunomodulator use were not eligible to participate in this study.

### Primary esophageal epithelial cell isolation

Esophageal biopsies were dissociated (27) and CD45+ cells were removed using positive isolation beads (STEMCELL TECHNOLOGIES). The untouched epithelial-enriched cell fraction was washed with PBS, lysed in Tri Reagent, then total RNA was extracted by column purification (PicoPure™ RNA Isolation Kit, Thermo Fisher Scientific).

### Esophageal epithelial cell culture

Telomerase-immortalized human esophageal keratinocytes EPC2-hTERT(28) were grown in keratinocyte-SFM (KSFM) containing 0.09mM Ca2+, 1ng/ml rEGF,1ng/ml bovine pituitary extract and 10,000 U/mL Penicillin-Streptomycin. High calcium KSFM media (1.8 mM Ca++) was used to induce epithelial maturation for 48 hours (i.e., flow cytometry, apoptosis assay, and MTT assay) prior to stimulation with cytokines. IFN-α (Fisher Scientific) and IFN-γ (Sigma-Aldrich) were added to culture media in varying concentrations, as indicated.

### Flow cytometry

EPC2-hTERT cells were dissociated, resuspended, then stained for viability (Zombie NIR^TM^, Biolegend) in PBS for 15min in FC Block (BD Biosciences). 1x10^6^ cells then stained in 200 µL on ice for 30 minutes with: Alexa Fluor® 405 anti-IFNα/β R1 (1µg, FAB245V, R&D Systems), PE anti-IFNα R2 (1µg, AB_2898715, Invitrogen), Alexa Fluor® 488 anti-IFNγ R1/CD119 (1µg, FAB6731G, R&D Systems) and APC anti-IFNγ R2 (0.5µg, FAB773A, R&D Systems). LSR-Fortessa and FlowJo software (BD Biosciences) were used for analysis.

### Western Blot

8 µg of protein was resolved on 10% SDS-PAGE, then transferred to a 0.2μm PVDF, blocked then incubated overnight at 4°C in primary antibody (rabbit anti-human STAT1 1:500, HPA000931, Sigma-Aldrich; mouse anti-human pSTAT1 1:1000, 33-3400, Invitrogen; mouse anti-human STAT2, 1:2500, MAB1666, clone 545117, R&D; rabbit anti-human pSTAT2, 1:1250, PA5-78181, Invitrogen). Membranes were washed then incubated for 1 hour at room temperature with secondary antibody (Goat Anti-Mouse Starbright Blue 520 and Goat Anti-Rabbit Starbright Blue 700, each at 1:2500, Bio-Rad). Blots were analyzed (ChemiDoc^TM^ MP Imaging System, Bio-Rad) and quantified using Image Lab software.

### 3D organoid cell culture

Esophageal organoids were cultured as described (27, 29). Briefly, 2 x 10^3^ EPC2-hTERT cells were mixed in 50µl of Matrigel (Corning®) and seeded into a 24-well plate, then cultured 500µl of high Ca^2+^ KSFM 11 days. Organoids were treated with IFN-α (Fisher Scientific) or IFN-γ (Sigma-Aldrich) on days 7 and 9. Organoid number, size, and formation rate (OFR) were determined from phase contrast images of organoids (BZ-X710 Fluorescence Microscope, Keyence). OFR was defined as the average number of ≥50 μm spherical structures at Day 11 divided by the total number of cells seeded in each well at day 0 (29). Organoids were isolated for downstream analysis using vigorous pipetting and vortexing in DPBS (Gibco) to dissociate them from the Matrigel matrix.

Organoids were paraffin-embedded (32), sectioned, and stained for hematoxylin and eosin (H&E) or immunohistochemistry (IHC) analyses. Basaloid cell content was determined from H&E stained organoids by calculating the total height of both basaloid cell layers divided by overall organoid diameter and expressed as a percentage (29). The average basaloid content was calculated using four measures per organoid. For IHC, organoid sections were incubated with anti-human pSTAT1 (1:200, 9167L, Cell Signaling Technology), anti-human Ki67 (1:200, ab16667, Abcam), anti-human IVL (1:20000, I9018-0.2ML, Sigma-Aldrich), anti-human CLDN1 (1:500, LS-C415827, LS-BIO), and anti-human OCLN (1:200, ab216327, Abcam). Semi-quantitative analysis was performed in ImageJ software (NIH) using a minimum of 10 organoids per group.

For gene expression analysis, day 11 organoids treated with IFNα 200U/ml (Fisher Scientific), IFN-γ 5ng/ml (Sigma-Aldrich), and IL-13 10ng/ml (Sigma-Aldrich) were isolated, washed, and lysed in Tri Reagent. RNA was extracted for sequencing using RNeasy Mini Kit (Qiagen) according to the manufacturer’s protocol.

### Bulk RNA sequencing and data analysis

RNA libraries of esophageal epithelial tissue biopsies and organoids were prepared using SMART-Seq HT Ultra Low Input RNA kit (Clontech) and NEBNext Ultra II RNA Library Prep Kit for Illumina, respectively. Paired-end sequencing was performed on Illumina HiSeq 2500 and generated data was deposited in NCBI GEO (18, 19) under accession numbers GSE234973 and GSE234424. Sequencing quality was assessed with FastQC 0.14.1 (30) and MultiQC 1.10 (31). Trimmomatic 0.39 (32) trimmed low-quality reads and removed adapters, followed by pseudo-alignment and quantification against GRCh38.p13 using kallisto 0.46.0 (33). Low-abundance genes were filtered, and weighted trimmed mean of M-values (TMM) normalization was performed using edgeR 3.38.4 (34). Principal component analysis (PCA) was performed using base R and differential gene expression was analyzed using limma 3.52.4 (35). Facilitated Gene set enrichment analysis (GSEA) was performed with clusterProfiler 4.4.4 (36, 37) using MSigDB (38, 39). A list of 429 genes with 4-fold elevated expression in the esophagus (esophageal-specific genes) was obtained from the Human Protein Atlas (accessed on August 11, 2022) (40). Heatmaps and interaction network plots were created using gplots 3.1.3 (41) and Cytoscape 3.9.1 (42). Statistical plots and other data visualizations were generated using the R package ggplot2 3.4.0 (43), unless otherwise stated.

### Air-liquid interface (ALI) culture

Cells were seeded on a 0.4µm transwell (Corning®) in 300µl of low-calcium KSFM as previously described (44). Cultures were grown in high calcium media beginning day 3, then on day 7 upper chamber media was removed. IFN-α (Fisher Scientific) and IFN-γ (Sigma-Aldrich) treatments were performed from day 10 to 14.

### Transepithelial electrical resistance (TEER)

Net ALI membrane resistance for each sample was determined by measuring the sample resistance with a MillicellERS-2 Voltohmmeter (Merck Millipore) then subtracting the blank sample resistance reading. TEER values (Ω * cm2) were calculated as net resistance * membrane area (0.33cm2 for 24-well inserts, Millipore). Only samples with TEER > 200 Ω*cm2 on day 10 were used for stimulation experiments.

### FITC-Dextran Permeability Assay

70kDa fluorescein isothiocyanate-dextran (FITC dextran; 3mg/ml; Sigma-Aldrich) was added to apical chambers and then incubated at 37°C for 4 hours. Basolateral chamber media was transferred into a clean 96-well plate and fluorescence levels were determined to assess flux.

### Caspase activity assays

Epithelial cells were matured, treated with cytokines, then incubated with 100µl of the caspase loading solution (1:200) for 60min. Caspase-3 and caspase-8 activity were determined using a fluorometric multiplex activity assay kit (Abcam) according to the manufacturer’s instructions. Fluorescence was measured at Ex/Em=535/620nm for caspase-3 and Ex/Em=490/525nm for caspase-8.

### MTT assay

Epithelial cells (1×10^4^/well) were cultured in 96-well plates, matured in high calcium media, treated with cytokines, then incubated with 0.5mg/ml MTT solution at 37°C for 3 hours. Cells were spun down at 2500rpm for 5min, then resuspended in 100µL of DMSO while shaking for 15 minutes at 300rpm. The absorbance was measured at 562nm.

## RESULTS

### Esophageal epithelium expresses interferon response signature in EoE

To identify cell types contributing to the interferon gene signature (IGS) observed in EoE biopsy gene expression data, we analyzed phenotypic clusters (Figure S1) generated by Morgan et al (8), that used samples from 6 patients with active EoE and 4 in remission. We used IGS from our prior work to examine gene expression across all cell types (Figure S2), observing the highest number of differentially regulated ISGs in epithelium (Figure 1A) compared to other cells (Figure S3).

**Figure 1.**
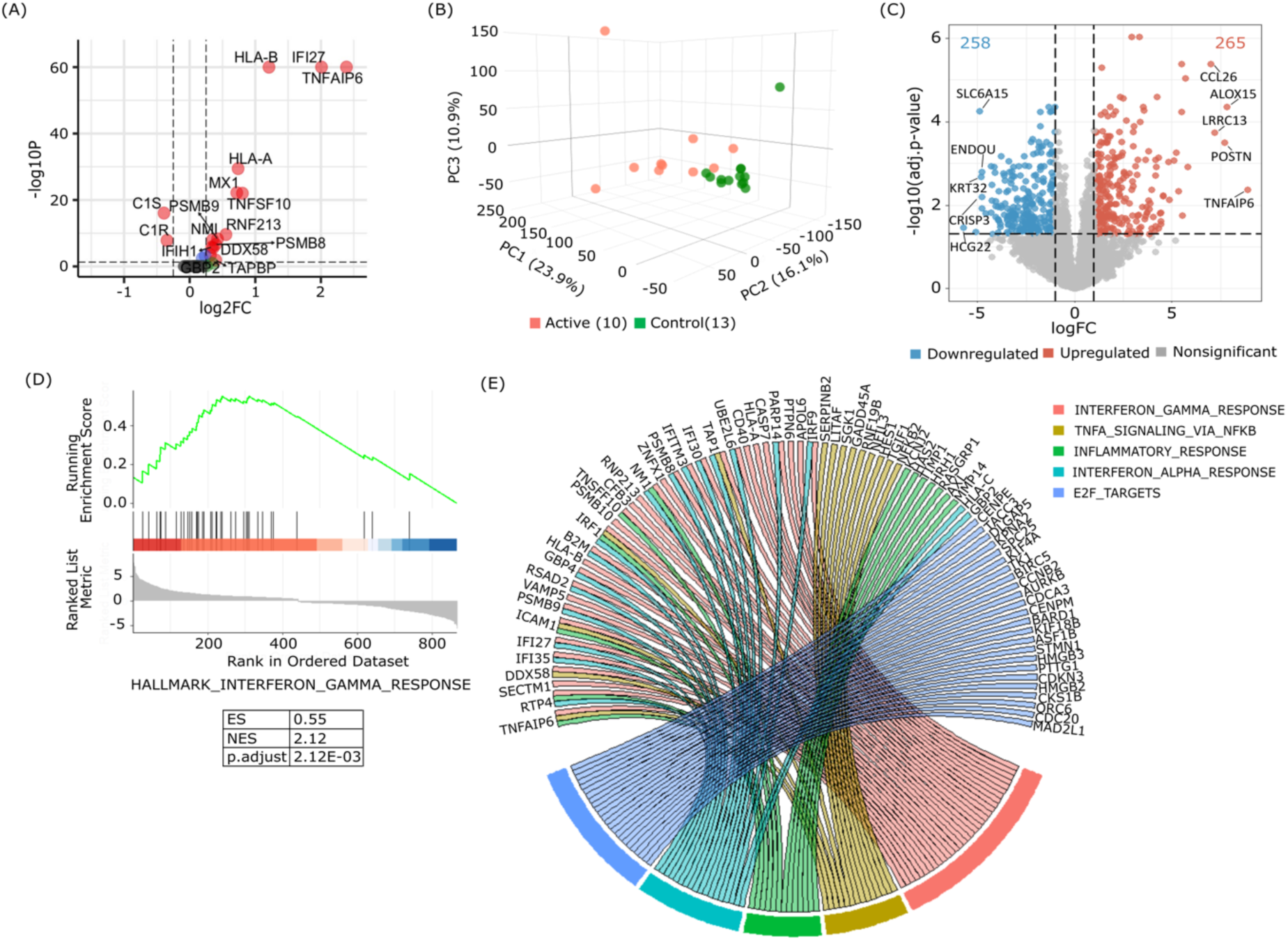
Transcriptomic profiling of esophageal epithelium in complementary datasets reveals enrichment of IFN response gene signature during active EoE. (A) Volcano plot of differentially expressed IFN signature genes between active and remission EoE in the esophageal epithelium single-cell cluster. (B) PCA plot showing separation in the comparison of EoE and control patients’ gene expression from CD45-depleted epithelial samples using the first three principal components that account for most of the variation in the dataset. (C) Volcano plot of DEGs in EoE active versus control, CD45-depleted, epithelial cell RNA-seq. (D) GSEA analysis revealed the Hallmark IFN-gamma response pathway as the most significantly enriched pathway in EoE epithelium. (E) Circos plot of overlapping differentially expressed genes in the top 5 significant enriched GSEA pathways from EoE compared to control.

To validate this finding, we isolated epithelial cells from patients with EoE and non-EoE controls (Table 1, patient demographics), and performed bulk RNA sequencing to analyze the epithelial-specific transcriptome in EoE. Analogous to findings in EoE whole-biopsy tissue (10, 12, 45, 46), unsupervised hierarchical clustering differentiates epithelial cell samples from active EoE biopsies from non-EoE controls (Figure S4 and Figure 1B). In epithelial-enriched samples from active EoE, we identified 523 differentially expressed genes (DEGs) compared to controls (FDR < 0.05 and log2FC ≥ Ι1Ι, Figure 1C and Table S2), with 265 upregulated and 258 downregulated. Upregulated DEGs included *CCL26* (47), *ALOX15* (48), *LRRC31* (49), *POSTN* (50), *TNFAIP6* (51), *KRT32* (52), and *CRISP3* (51), which have been previously implicated in the immunopathology of EoE.

**Table 1.** Demographics of patient cohort.

| Characteristics | ACTIVE (N = 10) | CONTROL (N = 13) |
| --- | --- | --- |
| Age (years) | 10 (4.5-12.0) | 14 (11-17) |
| Gender (male) | 8 (80%) | 7 (53.85%) |
| Eos/HPF | 87.5 (32.5-100) | 0 (0-0) |
| Swallowed Steroids (Y) | 3 (30%) | 0 (0%) |
| PPI (Y) | 4 (40%) | 9 (96.23%) |
| Diet (Restricted) | 8 (80%) | 5 (38.46%) |
| Race (White) | 5 (50%) | 13 (100%) |
| Ethnicity (Hispanic or Latino) | 2 (20%) | 0 (0%) |

Data are given as N (%) or median (interquartile range [IQR]) unless otherwise stated. IQR, interquartile range; HPF, high powered field; PPI, proton pump inhibitor.

Gene set enrichment analysis identified *IFN-γ response* (NES = 2.12, p.adjust = 2.12E-03) as the most significant upregulated pathway, followed by *TNF-α signaling by NFΚB* (NES = 1.93, p.adjust = 8.63E-03), inflammatory response (NES = 1.91, p.adjust = 8.63E-03), *IFNα response* (NES = 1.90, p.adjust = 1.32E-02), and a cell-cycle related pathway, *E2F targets* (NES = 1.93, p.adjust = 8.63E-03) (Figure 1D, Figure S5 and Table S3). A gene-pathway interaction network analysis (Figure 1E) indicated *TNFAIP6, RTP4, ICAM1, IRF1, NMI, and TAP1*, were shared across IFN-γ response, TNF-α signaling by NFΚB, inflammatory response, and IFN-α response pathways. There was a leading-edge overlap between the IFN-γ and IFN-α pathways, but no overlap genes in the E2F targets and other pathways.

### IFN-γ treatment reduces epithelial organoid formation and alters morphology

Given this evidence of pathway overlap, we next assessed the impact of interferon signaling on immortalized human esophageal keratinocytes, EPC2-hTERT (28). Flow cytometry confirmed interferon receptor expression in EPC2-hTERT (Figure 2A, gating strategy Figure S6). Western blot analysis demonstrated increase pSTAT-1 and pSTAT-2 following INF-γ and IFN-α treatment (Figure 2B), respectively, confirming functional receptor signaling.

**Figure 2.**
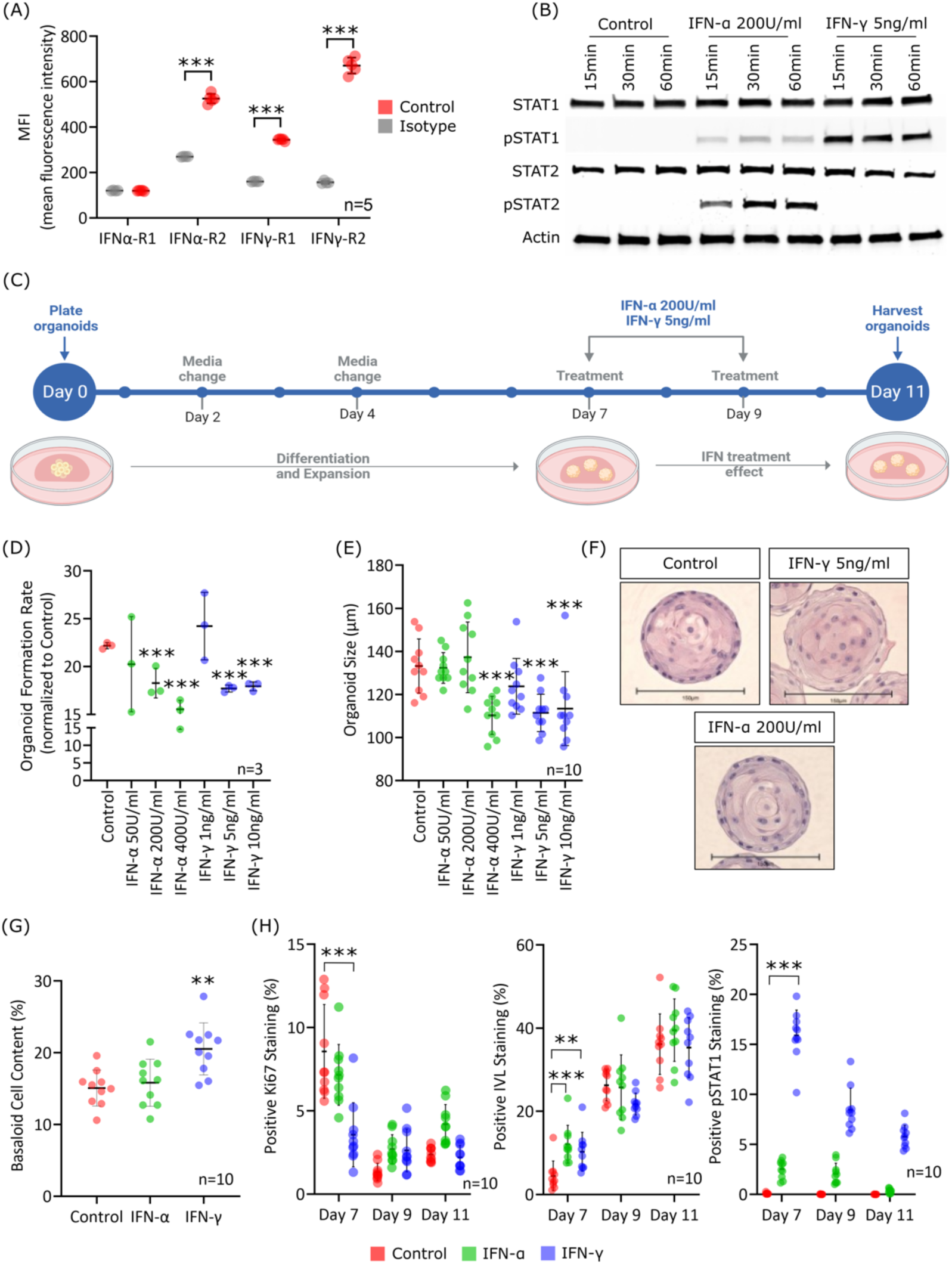
IFNγ-STAT1 stimulation induces changes in the morphological and functional features of human esophageal epithelial organoids. (A) Flow cytometry analysis of IFN-α and IFN-γ receptor subunits on EPC2-hTERT cells; plot showing the mean fluorescence intensity for each condition. n=5 replicates. (B) Time course of STAT1 and STAT2 phosphorylation by western blot in EPC2-hTERT cells, representative of 3 experiments. (C) Schematic depicting the 3D organoid epithelial culture model. The culture encompasses days 1 to 11, recapitulating the intricate architecture, proliferation, and differentiation gradient observed in the human esophageal epithelium. External stimuli are introduced into the media from days 7 to 9, capturing the responsive changes associated with esophageal disease conditions. (D) The organoid formation rate was determined on day 11 and calculated as the total number of organoids divided by the total number of seeded cells. (n=3 organoids per group, mean ± SD). (E) Organoid size was reported as the mean diameter (µm) at day 11 (n=10 organoids per group, mean ± SD). (F) Representative images of H&E staining of human esophageal epithelial 3D organoids harvested on day 11. Scale bar = 150 µm. (G) Basaloid content in organoids on day 11 was determined as the percentage of the total height of basaloid cell layers divided by the overall organoid diameter (n=10 organoids per group, mean ± SD, *** p-value < 0.001, ** p-value < 0.01, and * p-value < 0.05). (H) Immunohistochemistry staining for pSTAT-1, Ki67, and IVL in organoids treated with IFN-α or IFN-γ at day 11. The corresponding IHC images can be found in Figure S8. The graphs show the quantification of staining intensity (n=10 organoids per group, mean ± SD, *** p-value < 0.001, ** p-value < 0.01, and * p-value < 0.05).

Subsequently, we examined the effect of IFN-α and IFN-γ stimulation in the esophageal epithelial organoid model, due to its ability to reproduce the proliferation and differentiation gradient of the esophageal epithelium in humans (29). EPC2-hTERT esophageal keratinocytes in organoid culture were treated with IFN-α or IFN-γ on days 7 and 9 (Figure 2C). We observed decreased organoid formation rate in IFN-α 200U/ml (18.27 ± 1.26 %, p-value < 0.01), IFN-α 400U/ml (15.53 ± 0.76 %, p-value < 0.001) as well as IFN-γ 5ng/ml (17.7 ± 0.29 %, p-value < 0.001) and IFN-γ 10ng/ml (17.93 ± 0.38%, p-value < 0.001) groups (Figure 2D). Similarly, organoid size was significantly reduced following treatment with IFN-α 400U/ml (110.27 ± 8.41 µm, p-value < 0.001) as well as IFN-γ 5ng/ml (111.44 ± 8.23 µm, p-value < 0.001) and IFN-γ 10ng/ml (113.47 ± 16.29 µm, p-value < 0.01) (Figure 2E). Both IFN-γ and IFN-α display dose-dependent loss of central epithelial differentiation (Figure S7) but this was more pronounced with IFN-γ 5ng/ml and 200U/ml concentrations, respectively (Figure 2F). Further analysis of organoids treated with IFN-α 200U/ml and IFN-γ 5ng/ml revealed increased basaloid content in IFN-γ treated organoids (20.54 ± 3.64 % vs. 15.08 ± 2.50, p-value < 0.01), suggesting accumulation of outer, less-differentiated basaloid cell layers (Figure 2G). Staining for Ki67 showed a decrease at day 7 in IFN-γ treated organoids (p < 0.001), correlating with the strongest signal in the expression of pSTAT1 in all IFN-treated organoids (p < 0.001, Figure S8 and Figure 2H). There was a modest increase in IVL staining at day 7 in both IFN-α (p < 0.001) and IFN-γ treated groups (p < 0.01), consistent with earlier maturation in the interferon-treated samples.

### IFN-γ treated organoids mimic EoE epithelial gene expression signature

We used RNA-sequencing to examine transcriptional responses of IFN-γ and IFN-α treated organoids. Unsupervised sample clustering differentiated IFN-γ treated organoids from IFN-α and control groups (Figure 3A). We identified 1448 downregulated genes and 1859 upregulated genes in IFN-γ treated organoids at FDR < 0.05 and log2FC ≥ Ι1Ι (Figure 3B). In contrast, we detected 30 DEGs in IFN-α treated organoids at FDR < 0.05 and log2FC ≥ Ι1Ι (Figure S9, and Table S4). IFN-γ treated organoids share 27 upregulated and 97 downregulated transcripts with epithelium from EoE patients (Figure S10), more closely mimicking EoE patients’ epithelial gene expression than IFN-α treated organoids. This holds true with more permissive FDR thresholds for additional analysis of the IFN-α associated gene signature.

**Figure 3.**
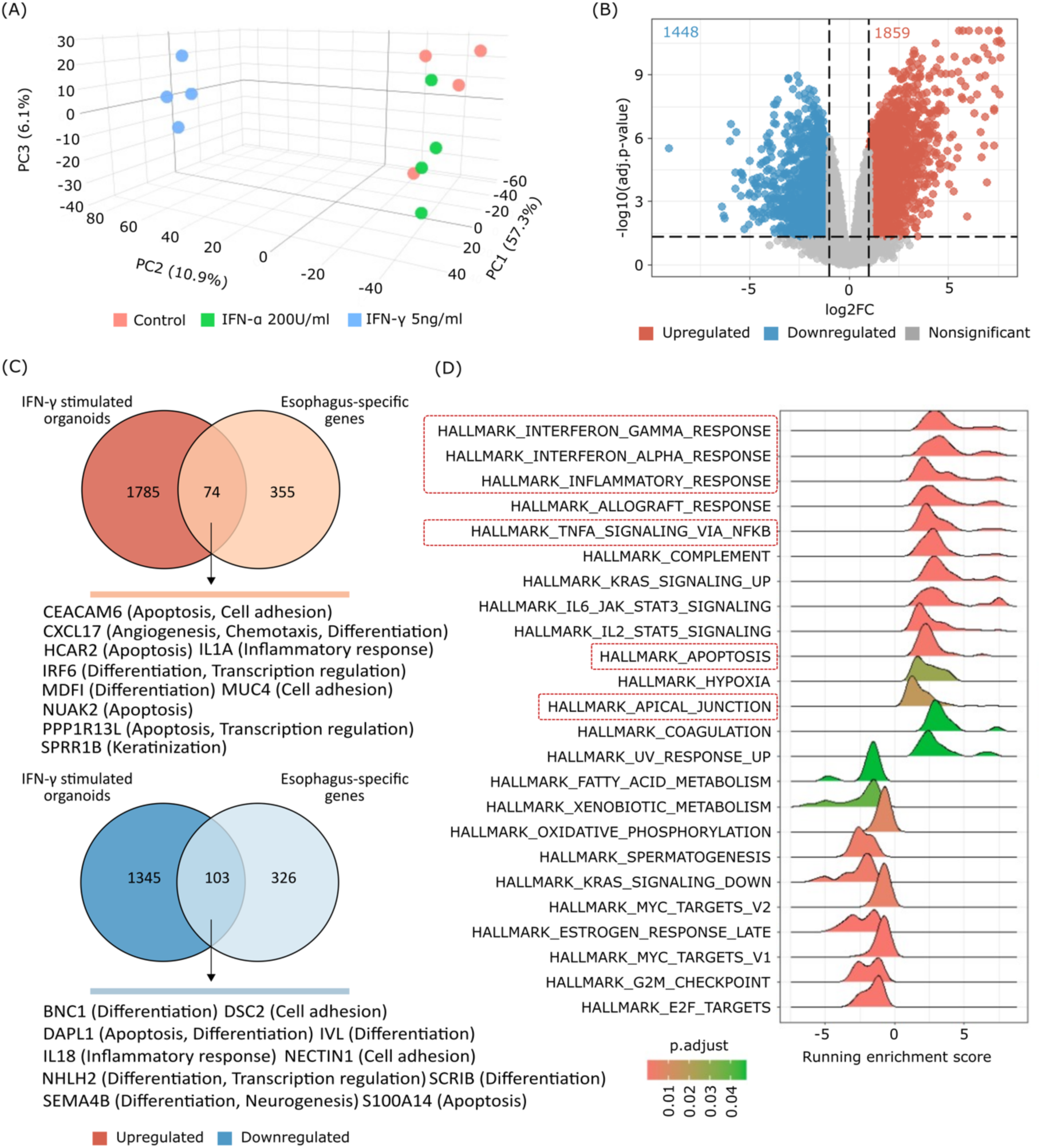
IFN-γ treatment *in vitro* replicates the *in vivo* IFN-γ response and changes the expression of a subset of genes relevant to esophageal biology. (A) PCA plot showing separation between IFN-γ treated organoids and IFN-α treated and unstimulated organoids based on gene expression profiles using the first three principal components that account for most of the variation in the dataset. (B) Volcano plot showing differential gene expression in IFN-γ treated organoids versus unstimulated organoid cultures. (C) The ridge plot shows the running enrichment score of 14 pathways with positive enrichment scores and 10 pathways with negative enrichment scores in IFN-γ treated organoids and p.adjust < 0.05. (D) Venn diagram illustrating the overlap of upregulated (red) and downregulated (blue) genes shared between esophagus-specific genes and IFN-γ treated organoids.

We investigated the impact of interferon signaling on esophageal-specific transcription using a dataset of esophagus-specific genes from the Human Protein Atlas (429 transcripts with >4-fold higher expression in the esophagus than other human tissues, Table S5) (53). We observed 74 (17.25%) upregulated and 103 (24%) downregulated DEGs overlapping between IFN-γ treated organoids with esophageal-specific genes (Figure 3D, Table S6). These genes are associated with a range of biological processes, including apoptosis, cell adhesion, and differentiation, suggesting IFN-γ treatment alters transcriptional regulation of pathways critical for epithelial cell proliferation, survival, and barrier function. We next tested the hypothesis that IFN-γ and IL-13 treatment result in distinct patterns of esophageal gene expression. We examined DEGs from IFN-γ treated and IL-13 treated organoids (54). Only 32 esophageal-specific genes are regulated by both IFN-γ and IL-13, implying that impacts of IFN-γ and IL-13 on esophageal-specific genes are largely distinct (Figure S11 and Table S7). Of these, 23 (71.8%) are downregulated. Gene set enrichment analysis of IFN-γ treated organoid DEGs identified 14 pathways with positive and 10 with negative enrichment scores (Figure 3D, Table S8)

### IFN-γ treatment disrupts the epithelial barrier in the esophageal epithelium

IFN-γ has been shown to contribute to epithelial barrier dysfunction within the intestine (55–60), and impaired barrier function is known to play a crucial role in EoE (61, 62). Our transcriptional analyses revealed positive enrichment of the HALLMARK_APICAL_JUNCTION gene set in IFN-γ treated organoids (Figure 3C, NES= 1.58, adj.p= 1.74E-02). Although 41 genes in this pathway are upregulated in the IFN-γ treated organoids (Figure 4A), there is decreased protein expression for claudin 1 and occludin in IFN-γ treated organoids (Figure 4B) consistent with prior reports of IFN-γ-induced tight junction disruption (56, 63, 64).

**Figure 4.**
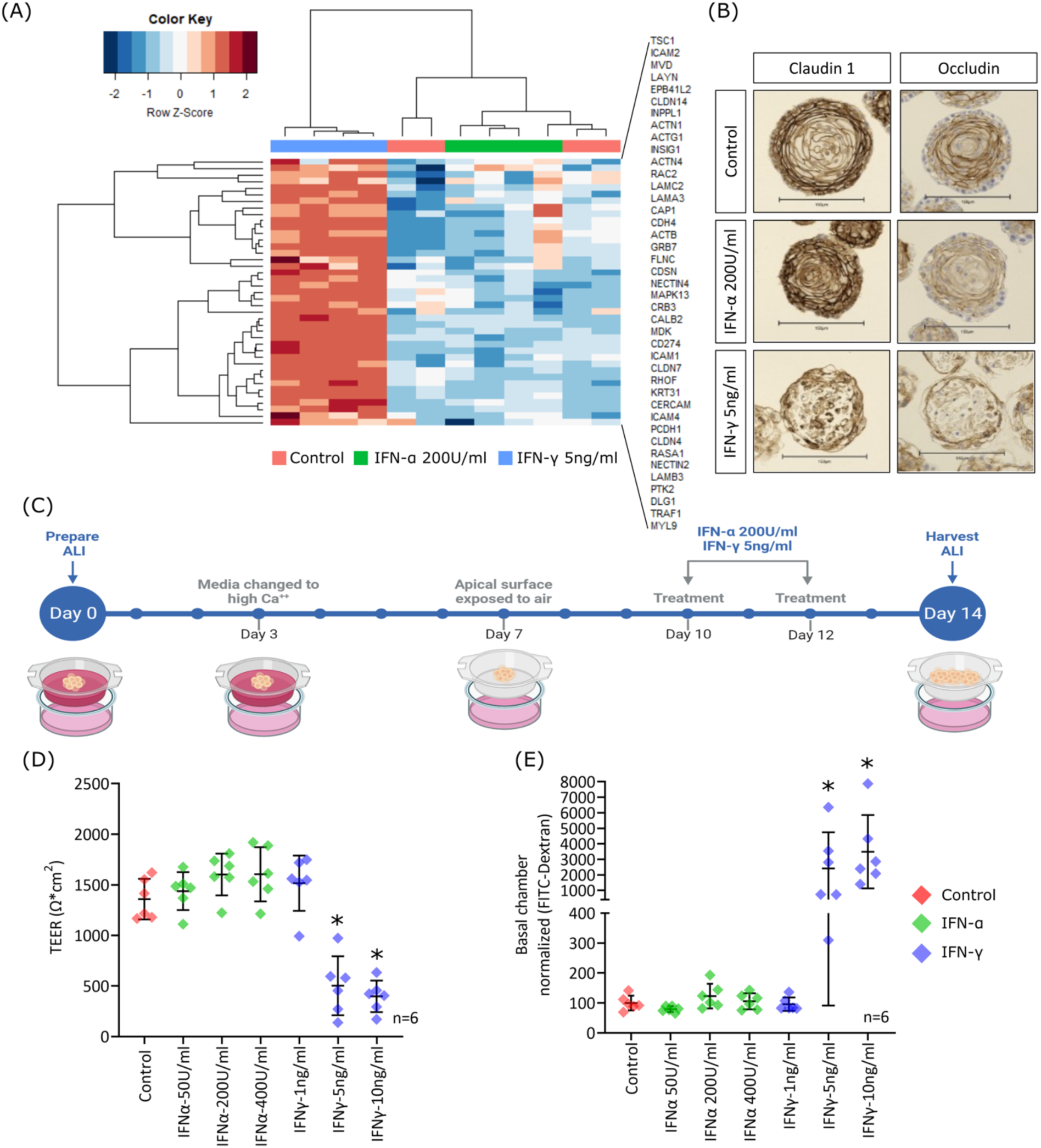
IFN-γ treatment disrupts epithelial barrier function in esophageal epithelium. (A) Hierarchical clustering based on the leading edge of the positively enriched pathway, HALLMARK_APICAL_JUNCTION; heatmap showing gene expression levels across IFN-γ and IFN-α treated and unstimulated organoids. (B) IHC of tight junction proteins, Claudin 1 and Occludin in 3D organoids derived from EPC2-hTERT cells treated with IFN-α and IFN-γ. (C) Schematic depicting the ALI (Air-Liquid Interface) epithelial culture model. The culture spans days 1 to 7, facilitating proliferation and initial differentiation, followed by days 7 to 10, which support terminal differentiation. External stimuli are introduced into the basolateral media chamber from days 10 to 14, enabling the evaluation of their impact on epithelial barrier function. (D)Transepithelial electrical resistance (TEER) was measured on day 14 and calculated as net resistance × membrane area (0.33cm2 for 24-well Millicell inserts), and only samples with TEER > 200 Ω*cm2 on day 10 were used for stimulation experiments. (n=6 wells per group, mean ± SD, * p-value < 0.05). (E) Basolateral translocation of 70kDa FITC-dextran across ALI membranes to assess the permeability upon IFN treatment (n=6 wells per group, mean ± SD, * p-value < 0.05).

We utilized the air-liquid interface culture (ALI) model (Figure 4C) to examine the effect of interferon treatment on esophageal epithelial barrier function. IFN-γ treatment decreased transepithelial electrical resistance (TEER) 3-fold in 5ng/ml and 10 ng/ml conditions (p-value < 0.05, Figure S12 and Figure 4D). Similarly, para-cellular permeability to 70 kDa FITC-Dextran increased after treatment with IFN-γ 5ng/ml and 10 ng/ml (p-value < 0.05 and <0.01, respectively, Figure 4E).

### IFN-γ treatment upregulates caspase expression in esophageal epithelium

The HALLMARK_APOPTOSIS gene set was positively enriched in IFN-γ treated organoids (Figure 3C, NES= 1.88, adj.p = 9.65E-04), and IFN-γ treated samples clustered separately from the other treatment conditions (Figure 5A), with upregulation of pro-apoptotic genes, including *PMAIP1, CASP7, TNFSF10, BCL2L1,* and *GADD45B*. To validate these findings, we examined caspase activity in EPC2-hTERT esophageal epithelial cells after 24 hours of IFN treatment in vitro. Caspase-3 and Caspase-8 activity were significantly increased in IFN-γ treated epithelial cells (Figure 5B, p-value < 0.001), accompanied by decreased cell viability with the highest concentration of IFN-α assessed (400U/ml, p-value < 0.001) and at all IFN-γ concentrations tested (p-value < 0.001). To assess the potential relevance of apoptosis in EoE, we stained esophageal biopsy tissue for activated caspase-3. We observed increased staining for activated caspase-3 in a cytoplasmic distribution scattered through the epithelium of EoE samples (Figure 5D).

**Figure 5.**
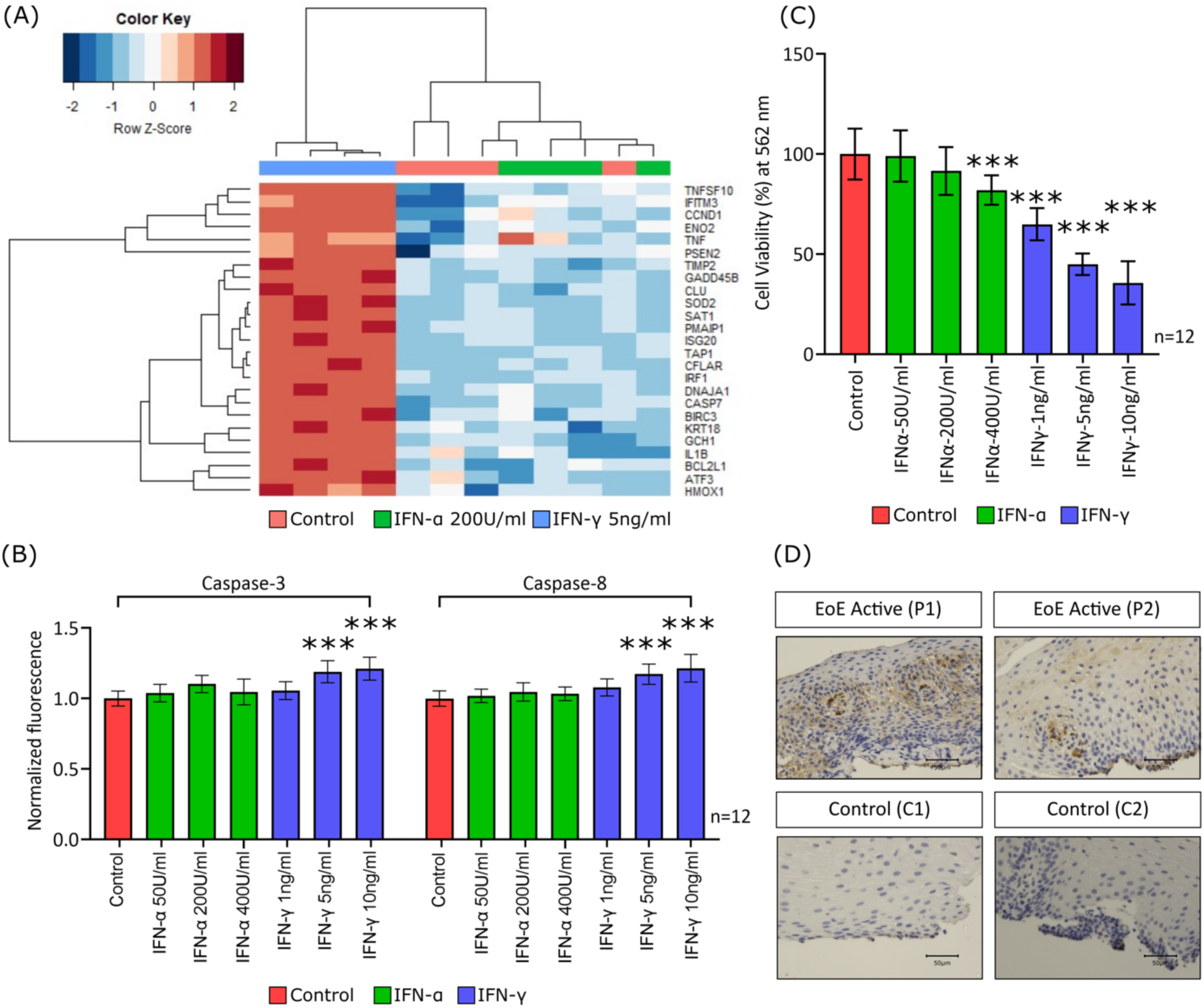
IFN-γ treatment potentiates cytotoxicity in esophageal epithelium. (A) Hierarchical clustering based on the leading edge of the positively enriched pathway, HALLMARK_APOPTOSIS; heatmap showing gene expression levels across IFN-γ and IFN-α treated and unstimulated organoids. (B) Expression of apoptotic markers, caspase-3 and caspase-8 in EPC2-hTERT cells treated with IFN-α and IFN-γ (n=12 replicates, mean ± SD, *** p-value < 0.001). (C) MTT assay to evaluate the cell viability upon IFN treatment (n=12 replicates, mean ± SD, *** p-value < 0.001). (D) Representative IHC for cleaved caspase-3 in the esophageal epithelium of EoE active and control subjects. Scale bar = 50 µm.

### Single-cell analysis reveals activated CD8+ T cells express *IFNG*

To determine which cells expressed interferons signaling genes, we examined the phenotypic clusters generated by Morgan et al. (8) as shown in Figure S1. Scaled expression frequency of *IFNG* in each cluster revealed enrichment in T cells (Figure 6A). In contrast, IFNA transcripts were not detected in the dataset, and IFNL transcripts were detected at low levels in <10% of all cells (data not shown). IFN-γ receptor subunits, *IFNGR1* and *IFNGR2*, were expressed by several cell types, with the highest expression observed in eosinophils and neutrophils (Figure 6A).

**Figure 6:**
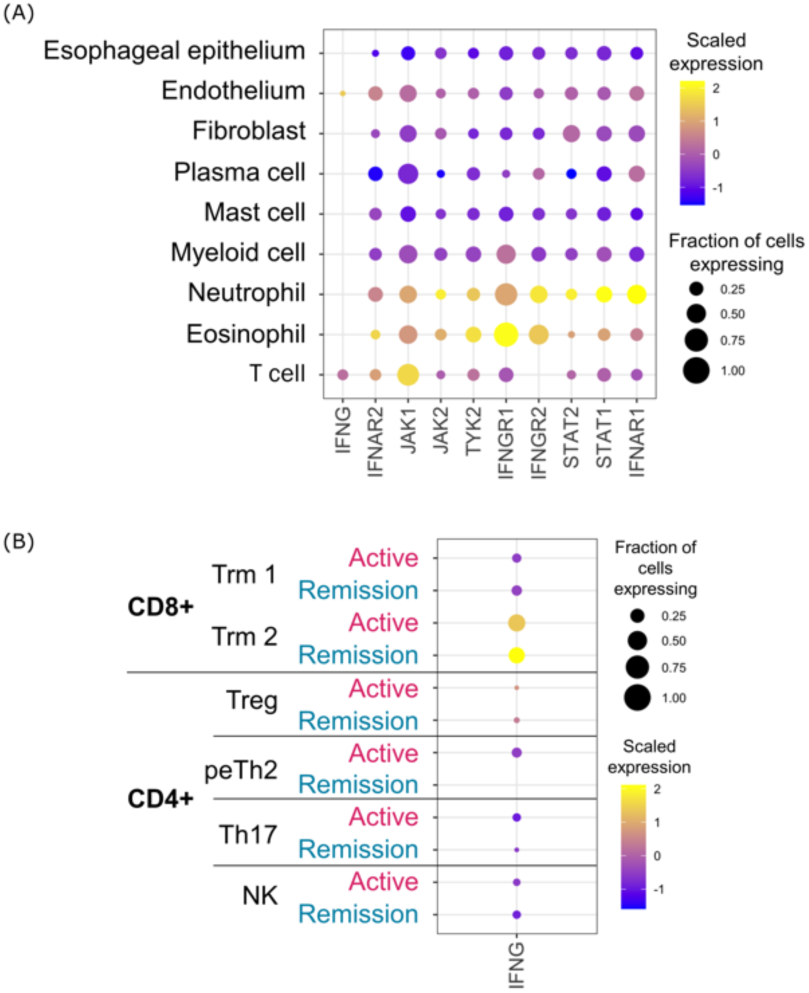
Single-cell analysis reveals activated CD8+ T cells as the main source of IFN-γ and upregulated IFN-γ receptors in eosinophils during active disease. (A) Dot plot of IFN signaling genes for each cell type in the esophagus displaying average expression and frequency of expression for each gene (B) Scaled expression and frequency of *IFNG* in esophageal T cells separated by disease state. The abbreviations used: Trm = tissue resident memory and peTh2 = pathogenic effector Th2.

To examine IFNG expression in T cells, we next interrogated T cell subsets using T cell cluster annotations as presented by Morgan et al. in Figure S13. This revealed that *IFNG* was expressed by a cluster of activated CD8+ T cells (Trm2) (Figure 6B). This cluster exhibited features consistent with activated tissue resident CD8+ memory cells, showcasing high enrichment in CD69 and FOS, and relative enrichment in CD103 (*ITGAE*) and CD49a (*ITGA1*) (Figure S13).

## DISCUSSION

In this study, we used complementary esophageal epithelial datasets to interrogate interferon signaling in EoE patients. We use scRNA-seq of EoE patients (6 with active EoE and 4 in remission) to identify that the epithelium has robust interferon signature gene (ISG) expression (Figure 1A), and to identify a CD8+ cell cluster with upregulated *IFNG* expression. RNA-sequencing analysis of esophageal epithelial cells isolated from biopsy tissue (Figures 1B-1E) confirmed ISGs as the most highly upregulated pathways. Using the esophageal organoid model, we demonstrate that IFN-γ treatment disrupts epithelial proliferation, decreases organoid formation and size, and causes abnormal morphology. IFN-γ treatment decreased epithelial barrier function in the air-liquid interface model and induced apoptosis *in vitro*.

Recent studies have highlighted that EoE is not exclusively a T2-driven disorder but has mixed pathophysiology (12). Esophageal epithelium has been shown to respond to multiple signaling mechanisms that are relevant in EoE, including cytokines (7, 9, 65), innate signaling molecules (44), and metabolites (54). Prior studies have identified elevated levels of IFN-γ expression in EoE patients (66, 67), *IFNG+* T cells in EoE mucosa (68). Prior studies in EoE have shown that ISG is upregulated and conserved in both children and adults with EoE (12, 45). In contrast, transcriptional studies in atopic dermatitis have revealed T2 and T17 signature among all age groups but T1 cytokine signatures primarily in adults (69, 70). Future studies that quantitate mucosal cytokine expression in EoE would help to determine a precise understanding of the balance of cytokine signals that impact esophageal epithelium during this disorder, leading to more sophisticated models and therapies.

Dysregulated IFN-γ signaling has been shown to contribute to increased intestinal permeability (14, 16, 17). These studies implicate IFN-γ-induced epithelial dysfunction via several mechanisms, which, to the best of our knowledge have not been examined in the esophageal epithelium. Our data demonstrate IFN-γ has detrimental effects on epithelial barrier function, as evidenced by decreased TEER and increased FITC-dextran permeability (Figure 4D and 4E). We observed a decrease in claudin 1 and occludin expression, consistent with prior reports of downregulation following IFN-γ exposure (64, 71). Further, we observe pronounced disruption of the typical proliferation timeline in IFN-γ treated organoid cultures, with decreased Ki67 on day 7, and significantly diminished OFR and size. Overall, this suggests that IFN-γ can have profound impact on esophageal epithelial function.

Our analysis of data from IFN-γ treated organoids indicated IFN-γ regulates 41% of esophageal-specific genes (Figure 3D). Classification based on biological processes revealed these affect apoptosis, cell adhesion, differentiation, and keratinization. IFN-γ has been shown to induce epithelial apoptosis in epithelial cells from other tissues (72–76), and we observed enrichment of apoptosis pathways IFN-γ treated organoid samples (Figure 5A). IFN-γ treatment induced caspase-3 and caspase-8 activity and cell death in esophageal epithelial cells. While increased epithelial apoptosis is not classically considered a finding of EoE, we observed increased cleaved caspase-3 staining in active EoE biopsy tissue samples. Skin lesions in atopic dermatitis, characterized by a mixed T2 high inflammatory infiltrate, have also been shown to have increased activated caspase-3 in keratinocytes(77, 78). Interestingly, in atopic dermatitis the degree of apoptosis has been correlated with dermal IFN-γ expressing CD4+ and CD8+ lymphocytes.

Prior studies have suggested that T cells may be a source of IFN-γ in the esophageal tissue of EoE patients. Wen *et al.* investigated 1088 individual T cells from EoE patients and control patients using scRNA-seq and reported *IFNG* expression in 84% of T cells (68). Sayej *et al*. reported that cultured biopsies from EoE patients had higher expression of TNF-α and IFN-γ, particularly in CD3+CD8+ but not in CD3+CD4+ T cell populations (11). Here, we identify *IFNG*-expression in a T cell cluster with characteristics consistent with CD8+ Trm given expression of CD8 subunits, *FOS*, CD103 (*ITGAE*), and CD49a (*ITGA1*) (79, 80). Our data indicate that these cells have similar level of *IFNG* expression in active and inactive EoE, suggesting they may maintain a more activated state even during inactive EoE. Apart from the CD8+ cluster, we observed modest expression of *IFNG* in the endothelium (Figure 6C). We did not detect meaningful levels of *IFNA* or *IFNL* in any cell clusters. It is possible that our gene expression studies failed to detect low levels of transcript expression that could be meaningful at the protein level. For example, recent data suggests the esophageal epithelium may produce IFN-γ *in vitro* (81). However, questions remain about role of interferon expression during EoE. Additional work to correlate the spatial organization of interferon expression in CD8+ cells with the epithelium could help determine if the ISGs signature in epithelium is related to proximity to CD8+ cells. The overall role of CD8+ cells during EoE is largely unknown, and examining these cells during active disease and remission may yield additional insight into the pathophysiology of EoE.

In summary, we provide evidence that esophageal epithelium in EoE patients expresses ISGs. Using organoid models, we demonstrate that IFN-γ signaling disrupts epithelial proliferation, barrier function, and induces apoptosis, indicating that IFN-γ has several detrimental effects on keratinocyte function in the esophagus which may have broad implications for infectious and inflammatory disorders of the esophagus. Our single cell analyses have identified a CD8+ *IFNG*-expressing Trm cell cluster present during active and inactive EoE as a potential source of IFN-γ in EoE. Additional work is needed to define the crosstalk between immune and epithelial cells to understand the extent to which interferon signaling may contribute to EoE pathogenesis.

### Patient and public involvement

Patients and/or the public were not involved in the design, or conduct, or reporting, or dissemination plans of this research.

## Author contributions

M.L., R.G., C.M.B., W.G.S., and M.A.R. planned and designed experiments. M.L, R.G., Z.M., T.T, J.B, M.C.C, C.M.B., and D.M.M performed experiments and discussed results. All authors contributed to the data analysis. A.B.M., W.E.S., M.A.R. were responsible for patient recruitment and study administration and support. M.L. wrote the first draft of the manuscript with input from M.A.R. All authors commented on and contributed to the final manuscript.

## Acknowledgments

We are grateful to the patients and their families for their participation in the studies. We acknowledge Azenta Life Sciences (Plainfield, NJ) for RNA sequencing of biopsies and esophageal epithelial organoid samples. We also acknowledge the University of Pennsylvania Perelman School of Medicine’s Center for Molecular Studies in Digestive and Liver Diseases (Molecular Pathology and Imaging Core (research Resource Identifier RRID: SCR_022420), as well as Children’s Hospital of Philadelphia Flow Cytometry and Pathology Core Facilities for technical support. We thank Lauren Dolinski and Tokunbo Ashorobi for their efforts in recruiting patients to participate in this study and Joshua X. Wang and Dr. Takeo Hara for valuable technical guidance regarding organoid culture. We also thank members of the Center for Pediatric Eosinophilic Disorders for helpful discussions.

