## Supplemental figures for "Interferon-γ signaling in eosinophilic esophagitis has implications for epithelial barrier function and programmed cell death": Supplemental_20240126_bioRxiv.pdf

### Human Interferon-γ Impairs Epithelial Barrier Function and Potentiates Cytotoxicity in Eosinophilic Esophagitis

#### Supplemental Figures

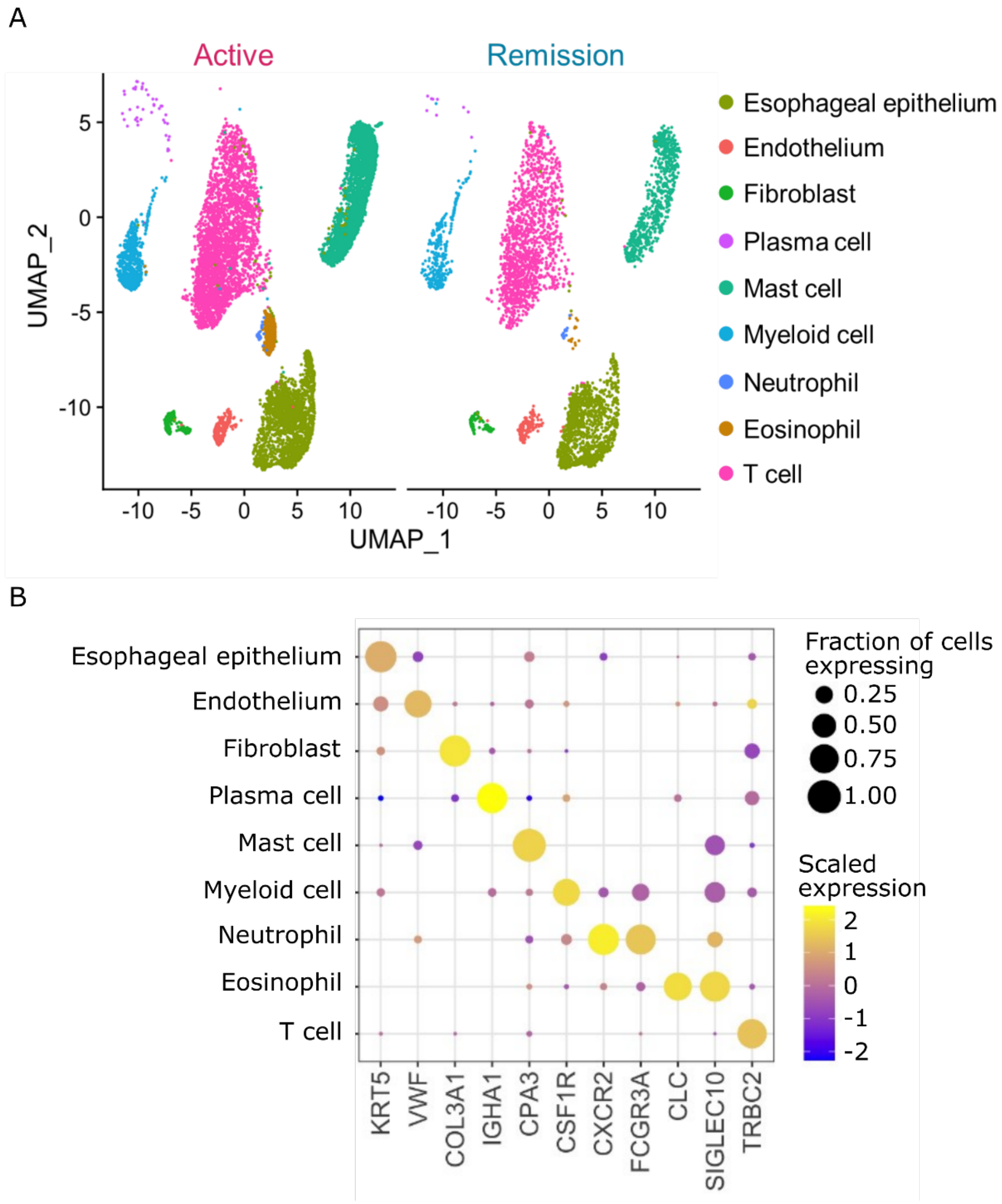


Supplemental Figure S1. (A) Uniform manifold approximation projection (UMAP) and (B) marker genes of clusters identified in the single-cell analysis of esophageal biopsy tissue from Morgan *et. al* (GSE175930) with *n=10,049 cells for active disease and 4,194 cells for remission.*

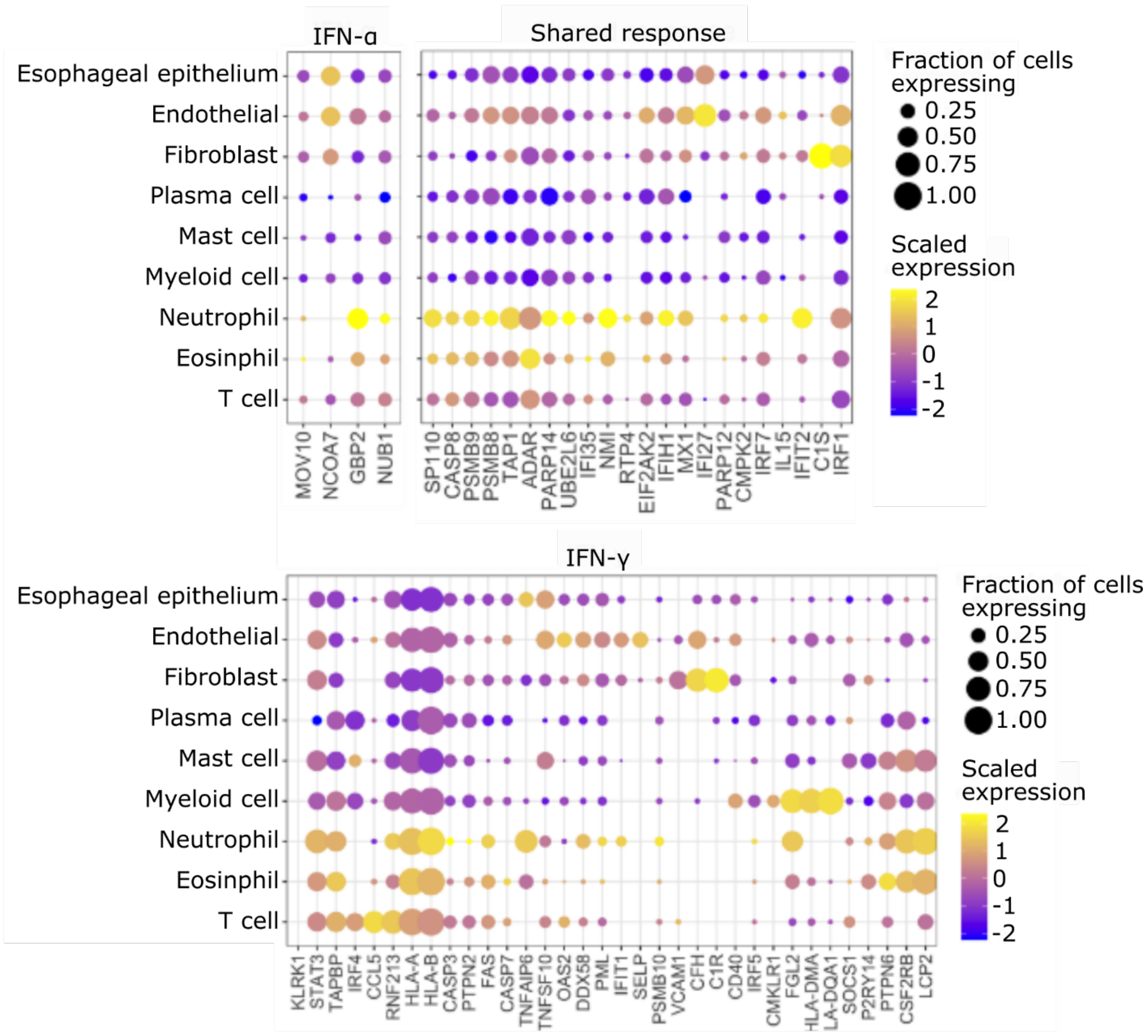


Supplemental Figure S2. Scaled expression and frequency of IFN- γ, IFN-α, and shared interferon signature genes from single cell analysis of active disease in the esophagus. Gene sets were identified between adult and pediatric EoE and control biopsy tissue in Ruffner et al, *J. Immunol.* 2021;206(6):1361–1371. *IFNA* and *IRF9* were not identified in this dataset, and *IFNL1* was identified in <10% of cells in each cluster.

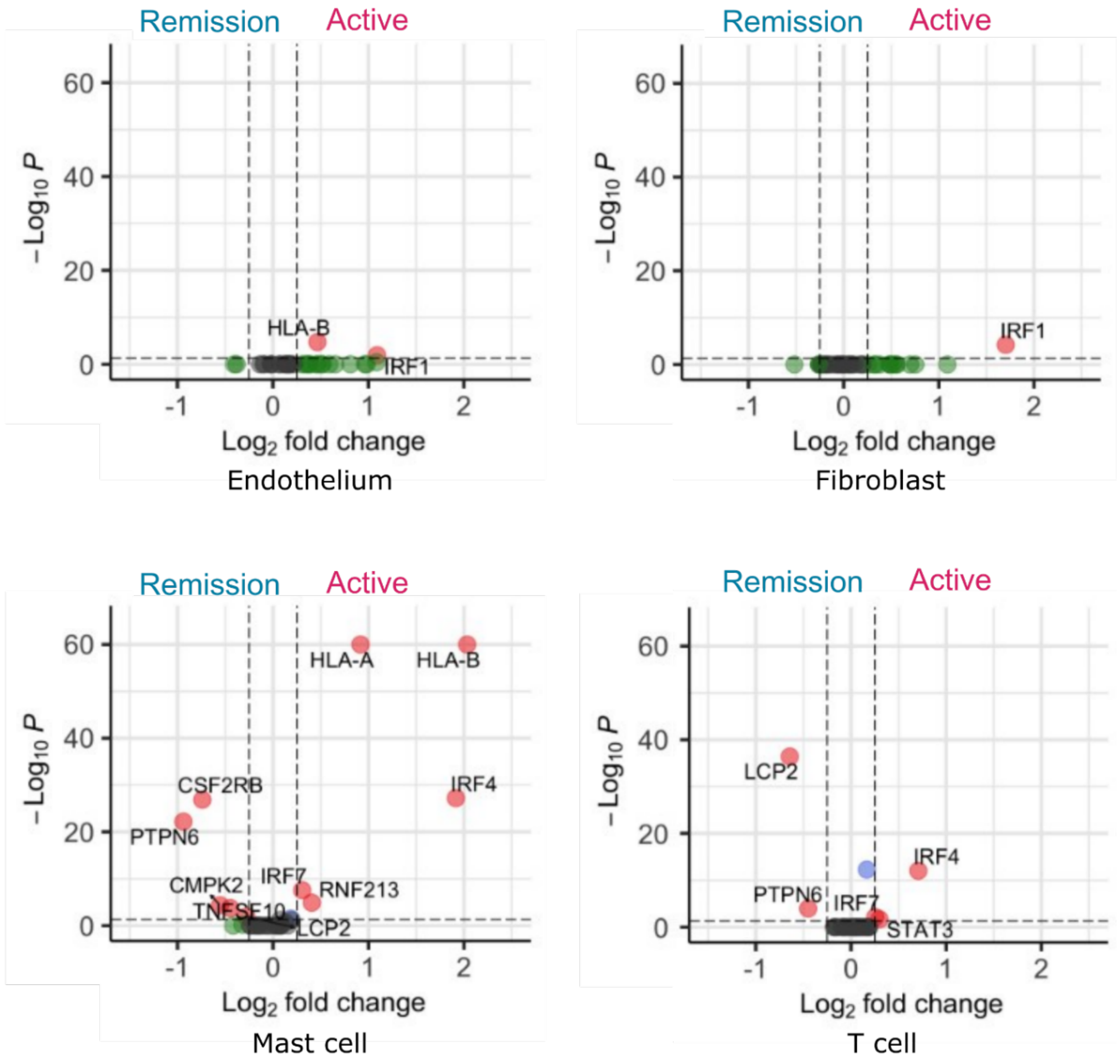


Supplemental Figure S3. Volcano plots displaying differentially expressed interferon signature genes between active and remission states in selected single-cell RNAseq cell clusters. The thresholds for differential expression were set at p-value <0.05 and -0.25 > log2FC > 0.25. Clusters without differential expression of interferon signature genes were excluded.

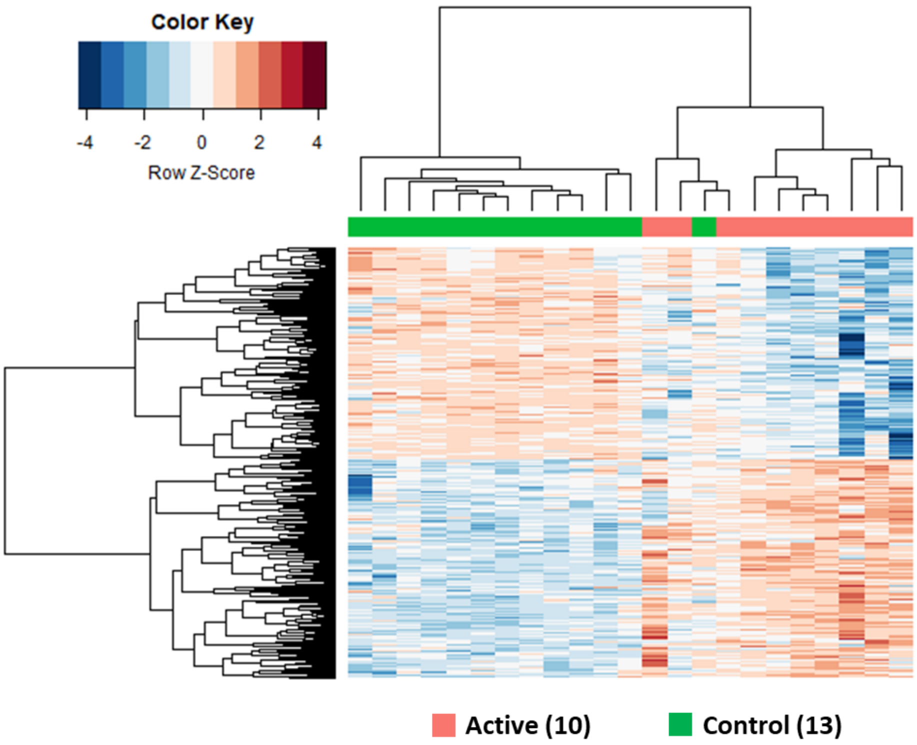


Supplemental Figure S4. Unsupervised hierarchical clustering of the expression levels of the most variable 523 genes in epithelial cell RNA-sequencing data from EoE active and control samples. The heatmap displays the expression levels of the genes with the highest variance across samples, with rows representing genes and columns representing samples.

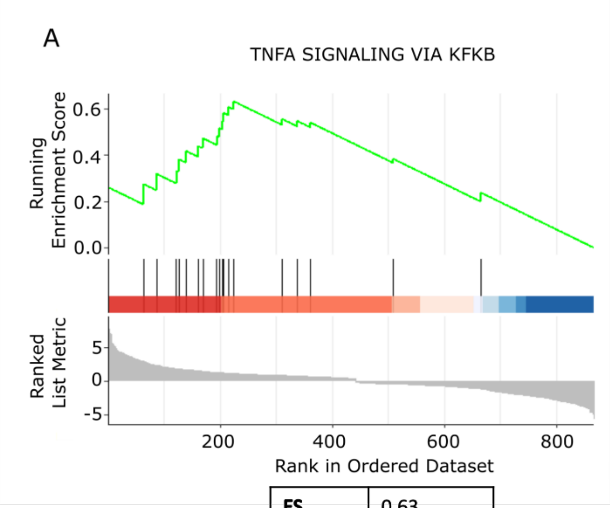


| | |
|---|---|
| **ES** | 0.63 |
| NES | 1.93 |
| p.adjust | 8.63E-03 |

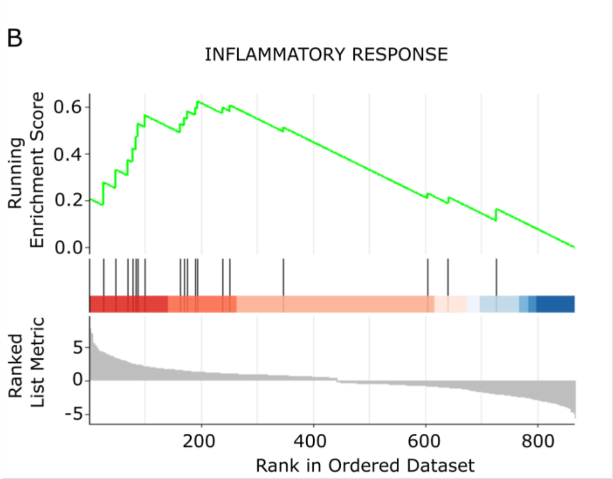


| | |
|---|---|
| **ES** | 0.62 |
| NES | 1.91 |
| p.adjust | 8.63E-03 |

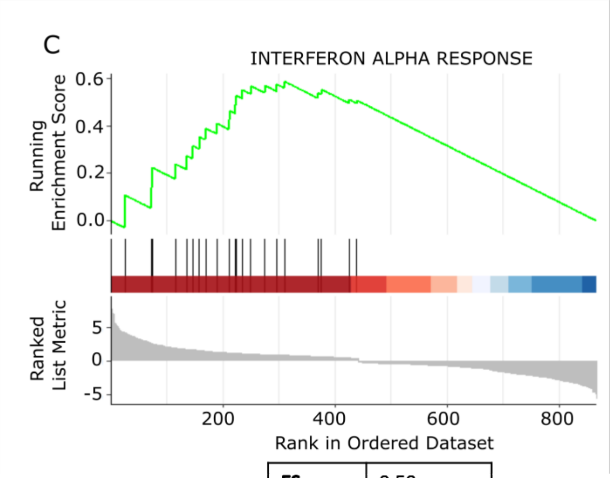


| | |
|---|---|
| **ES** | 0.59 |
| NES | 1.90 |
| p.adjust | 1.32E-02 |

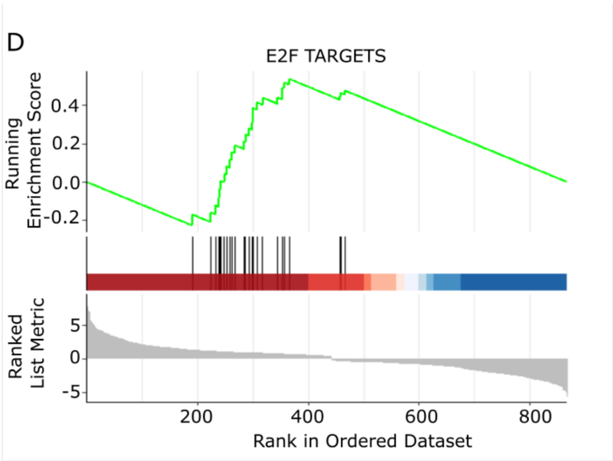


| | |
|---|---|
| **ES** | 0.54 |
| NES | 1.85 |
| p.adjust | 1.49E-02 |

Supplemental Figure S5. Gene set enrichment analysis (GSEA) of epithelial cell RNA-sequencing data from EoE active and control samples demonstrated enrichment for INTERFERON_GAMMA_RESPONSE (Figure 1D as the top enriched pathway, followed by A) TNFA_SIGNALING_VIA_NFKB, B) INFLAMMATORY_RESPONSE, C) INTERFERON_ALPHA_RESPONSE and D) E2F_TARGETS.)

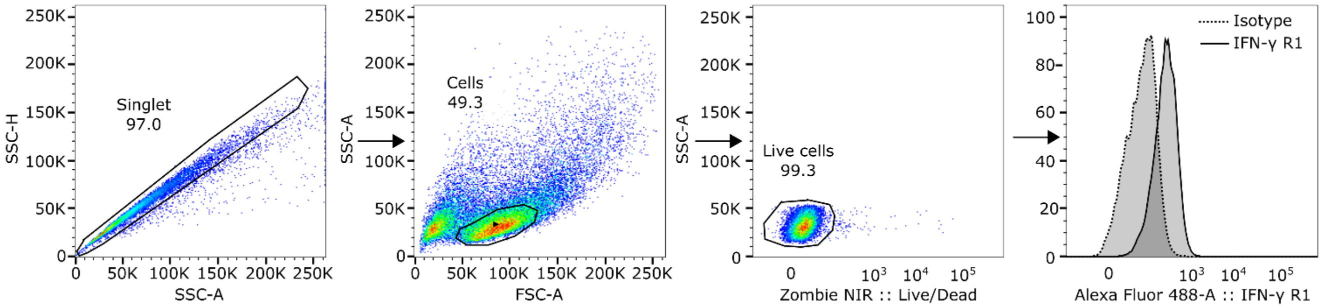


Supplemental Figure S6: Gating schematic for experiments assessing expression of interferon receptors on EPC2-hTERT esophageal epithelial keratinocytes. Cells were gated using FSC and SSC to identify single cells, dead cells were excluded using Zombie amine-reactive live/dead dye, then fluorescence was identified and compared to isotype control-stained cells.

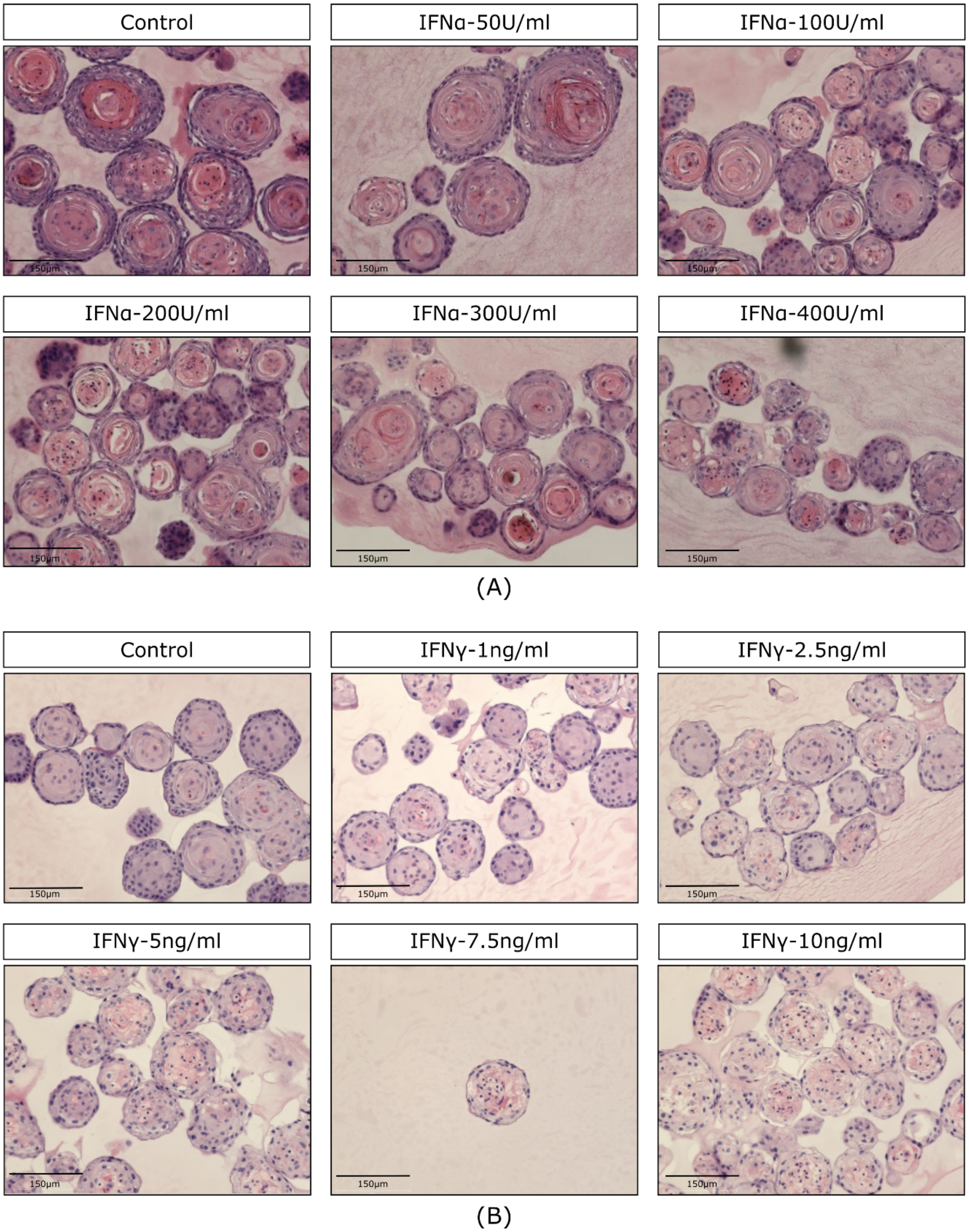


Supplemental Figure S7: EPC2-hTERT esophageal epithelial organoids were harvested at day 11, fixed embedded, and stained with hematoxylin and eosin to assess the impact of interferon treatment on organoid architecture. (A) At and above concentrations of 200U/ml, IFN- α organoids show disrupted central stratification. (B) At and above concentrations of 5ng/ml, IFN-γ treated organoids show disruption in central stratification. Scale bar = 150 μm.

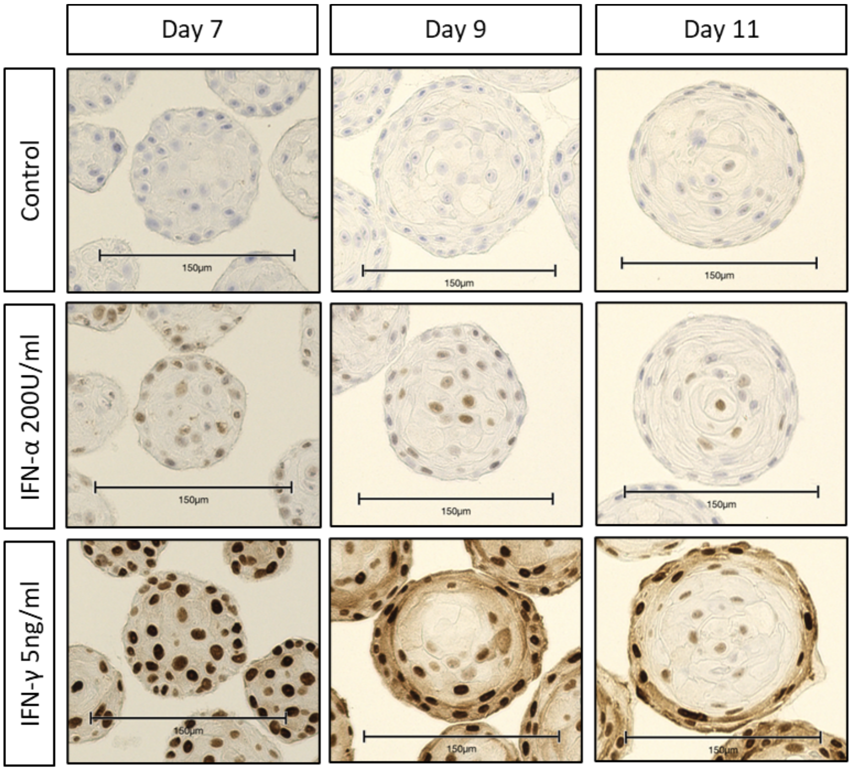

(A) pSTAT-1

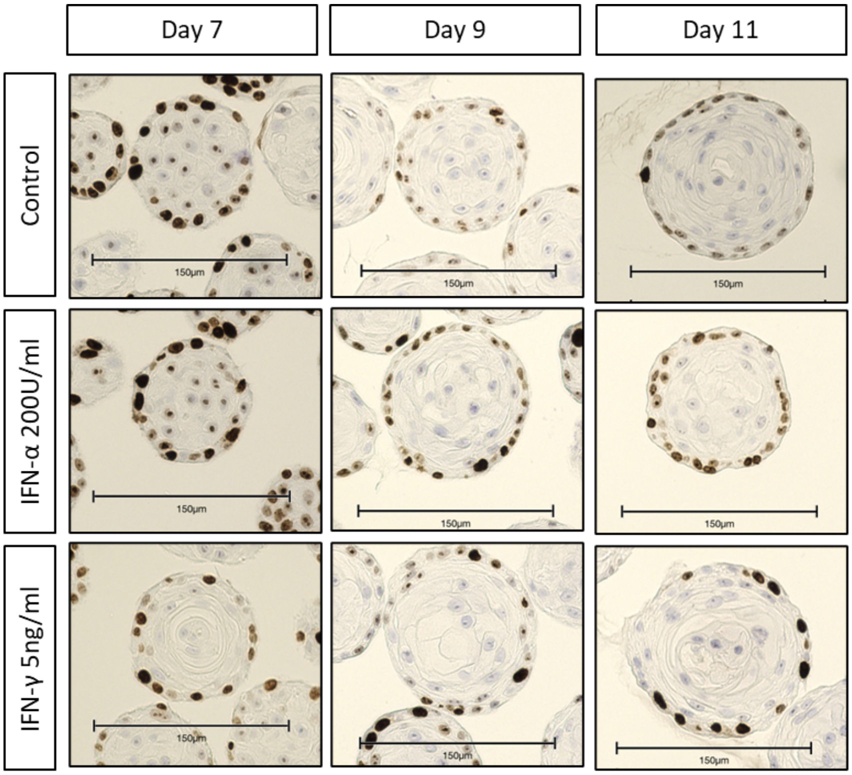

(B) Ki-67

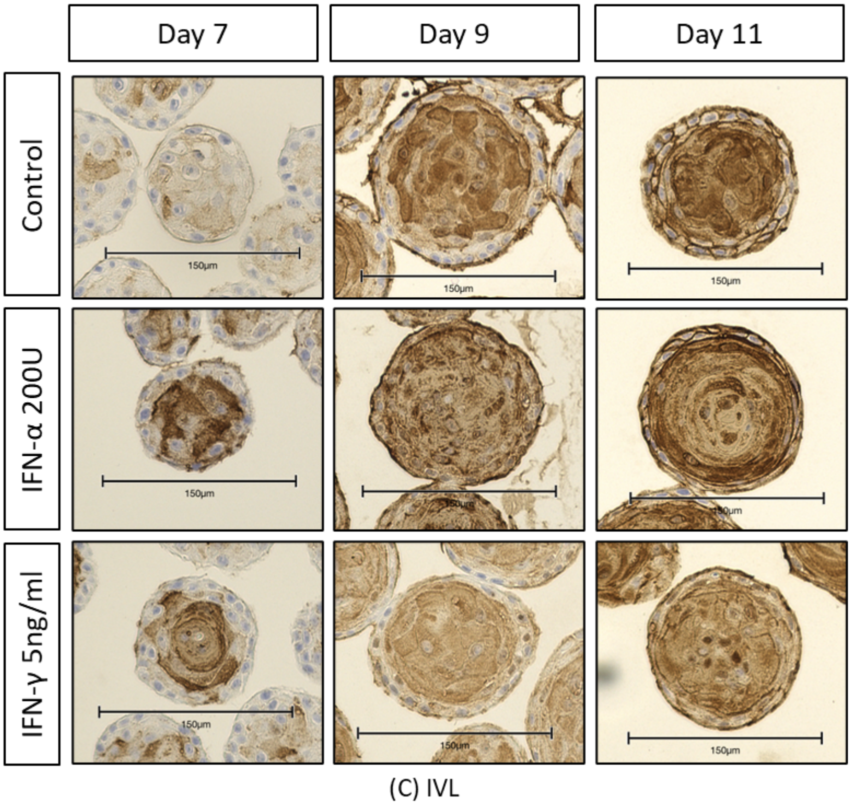


Supplemental Figure S8. Morphological characteristics of human esophageal epithelial 3D organoids treated with IFN-α 200U/ml and IFN-γ 5ng/ml. The organoids were harvested on Day 7, Day 9, and Day 11 and stained for (A) pSTAT-1 to assess for IFN signaling, (B) Ki67 to assess proliferation, and (C) involucrin (IVL) to assess epithelial differentiation. Scale bar = 150 µm.

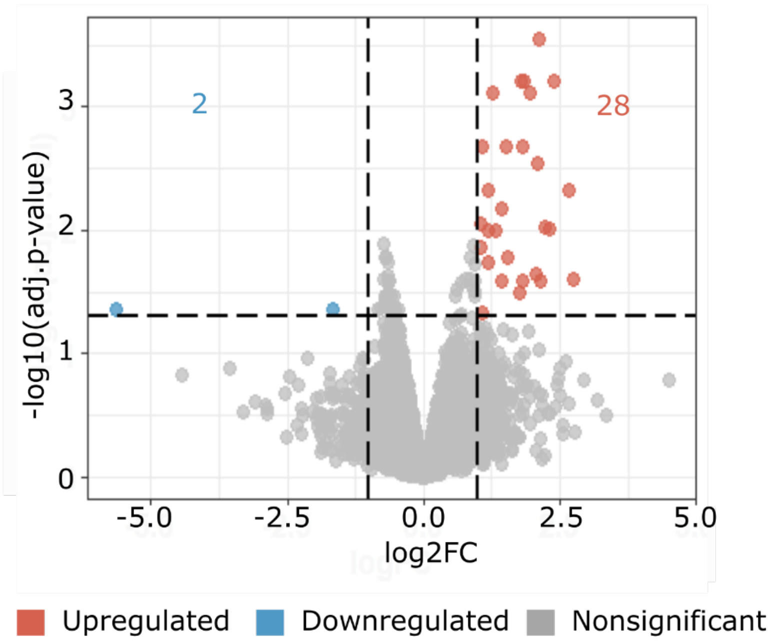


Supplemental Figure S9. Volcano plot showing differentially expressed genes in IFN-α treated organoids versus unstimulated organoid culture at the threshold FDR < 0.05 and log2FC ≥ │1│

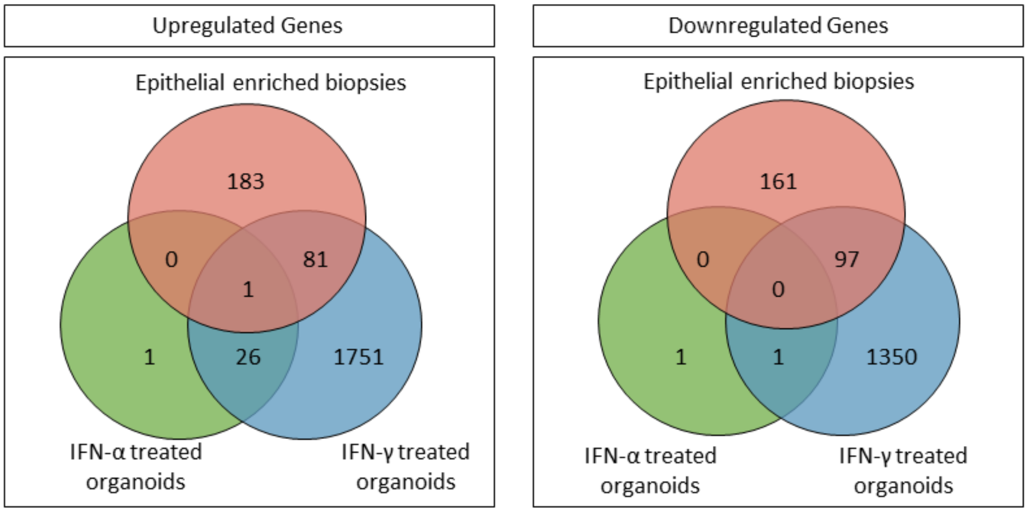


Supplementary Figure S10. Venn diagram illustrating the overlap of upregulated and downregulated genes shared between EoE epithelial cells (active vs. control), IFN-α (green) and IFN-γ (blue) treated organoids at FDR < 0.05 and log2FC ≥ │1│

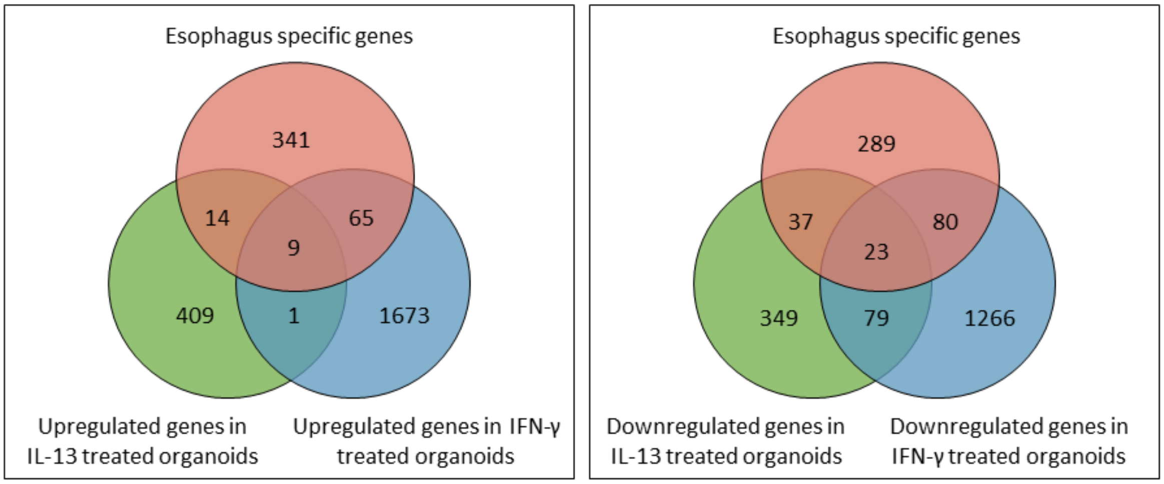


Supplementary Figure S11. Venn diagram illustrating the overlap of upregulated and downregulated genes in IL-13 (green) obtained from Hara T, et al, Cell Mol Gastroenterol Hepatol. 2022;13(5):1449–1467and IFN-γ (blue) treated organoids is compared to 429 esophagus-specific genes (orange). The esophagus-specific gene list was derived from the Human Protein Atlas. It was determined as the list of transcripts expressed 4-fold higher specifically in the esophagus when compared to other tissue types. This analysis was performed to assess how many esophagus-specific genes are affected by each type of cytokine treatment.

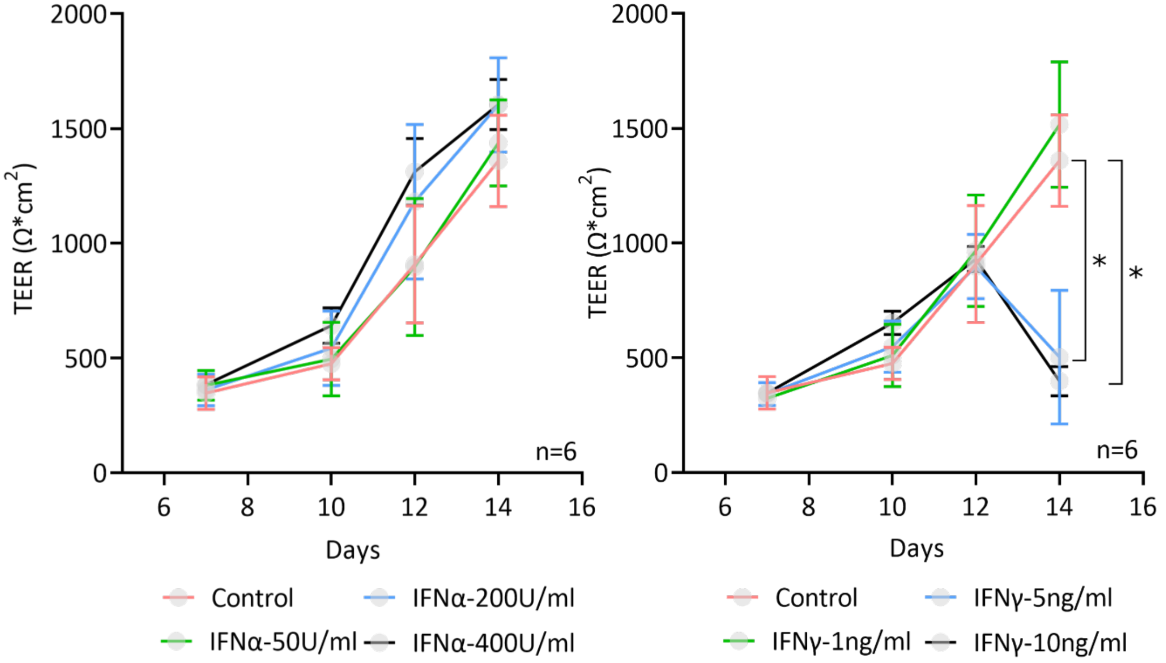


Supplemental Figure S12. EPC2-hTERT cells in air-liquid interface culture were treated with IFN-α and IFN-γ from days 10-14. Transepithelial electrical resistance (TEER) was measured and calculated as net resistance × membrane area (0.33cm2 for 24-well Millicell inserts). (n=6 wells per group, mean ± SD, * p-value < 0.05).

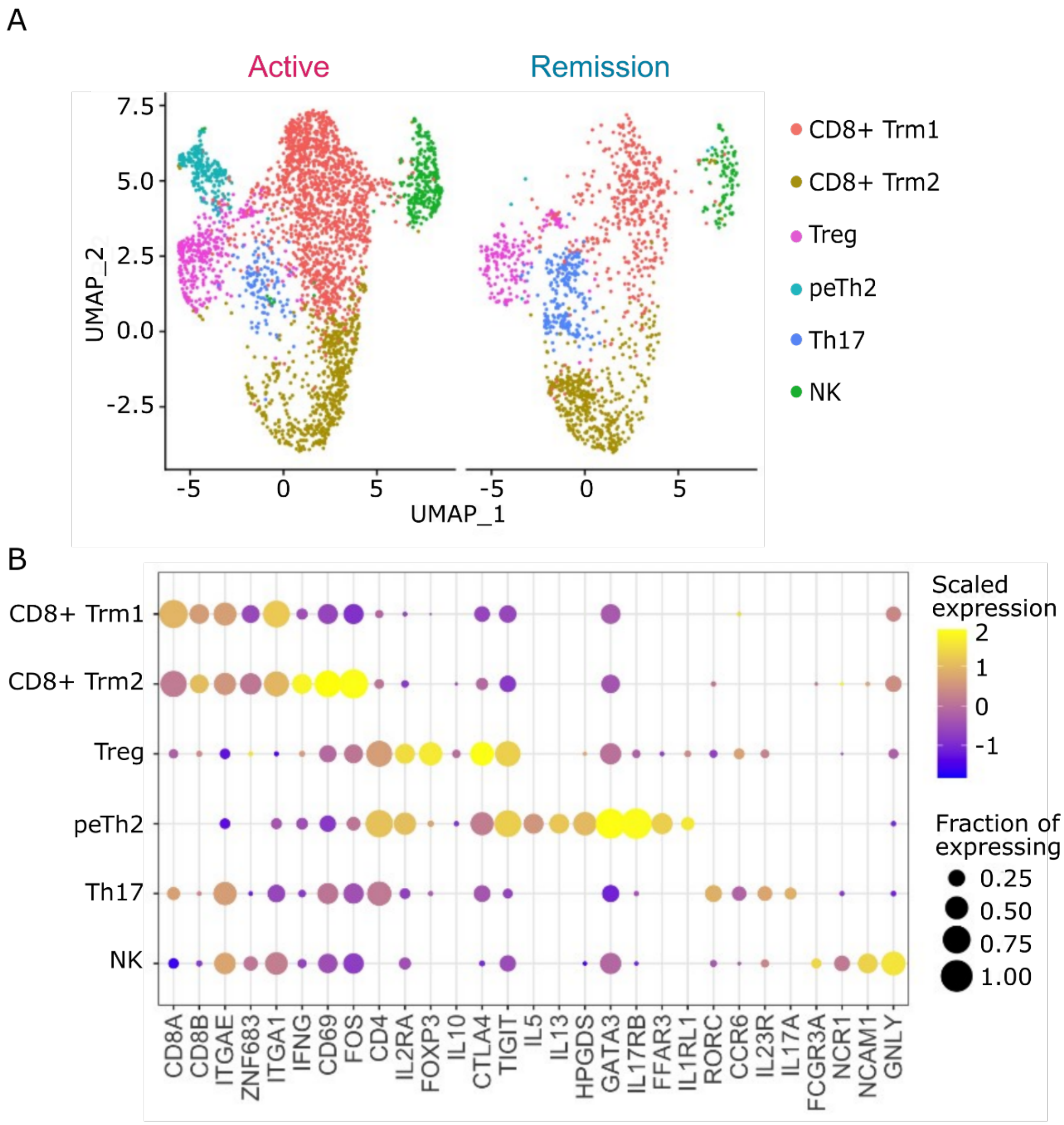


Supplemental Figure S13. (A) Uniform manifold approximation projection (UMAP) and (B) marker genes of clusters identified in single-cell data of T cells obtained from esophageal biopsy tissue. The clusters are identified based on the expression of marker genes, with a total of 3067 cells for active disease and 1357 cells for remission analyzed. The table included presents the percentage of cells in each cluster. The abbreviations used: Trm = tissue resident memory and peTh2 = pathogenic effector Th2.
